# Intrinsic circadian timekeeping properties of the thalamic lateral geniculate nucleus

**DOI:** 10.1101/2021.05.06.442920

**Authors:** L. Chrobok, K. Pradel, M.E. Janik, A.M. Sanetra, M. Bubka, J. Myung, A.R. Rahim, J.D. Klich, J.S. Jeczmien-Lazur, K. Palus-Chramiec, M.H. Lewandowski

**Affiliations:** Department of Neurophysiology and Chronobiology, Institute of Zoology and Biomedical Research, Jagiellonian University in Krakow, Gronostajowa Street 9, 30-387 Krakow, Poland; Department of Glycoconjugate Biochemistry, Institute of Zoology and Biomedical Research, Jagiellonian University in Krakow, Gronostajowa Street 9, 30-387 Krakow, Poland; Graduate Institute of Mind, Brain, and Consciousness, Taipei Medical University, 172-1 Sec. 2 Keelung Road, Da’an District, Taipei 106, Taiwan; Brain and Consciousness Research Centre, Taipei Medical University-Shuang Ho Hospital, Ministry of Health and Welfare, 291 Zhongzheng Road, Zhonghe District, New Taipei City 235, Taiwan

**Keywords:** circadian clock, clock genes, lateral geniculate nucleus, light-entrainable oscillator, multi-channel electrophysiology, PER2::LUC bioluminescence

## Abstract

Circadian rhythmicity in mammals is sustained by the central brain clock – the suprachiasmatic nucleus of the hypothalamus (SCN), entrained to the ambient light-dark conditions through a dense retinal input. However, recent discoveries of autonomous clock gene expression cast doubt on the supremacy of the SCN and suggest circadian timekeeping mechanisms devolve to local brain clocks. Here we use a combination of molecular, electrophysiological and optogenetic tools to evaluate intrinsic clock properties of the main retinorecipient thalamic centre – the lateral geniculate nucleus (LGN). We identify the dorsolateral geniculate nucleus (DLG) as a slave oscillator, which exhibits core clock gene expression exclusively *in vivo*. Additionally, we provide compelling evidence for intrinsic clock gene expression accompanied by circadian variation in neuronal activity in the intergeniculate leaflet (IGL) and ventrolateral geniculate nucleus (VLG). Finally, our optogenetic experiments propose the VLG as a light-entrainable oscillator, whose phase may be advanced by retinal input at the beginning of the projected night. Altogether, this study for the first time demonstrates autonomous timekeeping mechanisms shaping circadian physiology of the LGN.

## INTRODUCTION

Benefits and risks of the outside world undergo dramatic daily changes governed by the light-dark cycle. Thus most, if not all, living organisms developed biological clocks, which enable them not only to react to these cyclic events but also to predict them. In mammals, the central clock is localised in the suprachiasmatic nuclei (SCN) of the hypothalamus (Hastings *et al*., 2018, 2019). Its timekeeping properties are provided by clock cells maintaining a transcription-translation feedback loop (TTFL) of core clock genes whose expressions are regulated by their own protein products in the period of circa 24 h (Takahashi, 2017). Circadian rhythms on the molecular level are reflected in the daytime rise in the electrical activity of the SCN neurons followed by their nocturnal silencing (Belle *et al*., 2009; Colwell, 2011). Additionally, the SCN is best known for its photoentrainability, as it receives a dense innervation from the retinal ganglion cells (heavily from these synthesising a ‘circadian photopigment’ melanopsin) and responds to ambient light by both neurophysiological excitation and phase shifts in clock gene expression (Beier *et al*., 2020).

The lateral geniculate nucleus (LGN) of the thalamus poses one of the main gateways for photic information conveyed from the retina to the rest of the brain. Specifically, the LGN forms a complex of three independent retinorecipient structures: (1) the dorsolateral geniculate nucleus (DLG), with its thalamo-cortical neurons reaching the primary visual cortex and thus being directly involved in image-forming vision (Sherman, 2005), (2) the intergeniculate leaflet (IGL) implicated in circadian photoentrainment (Albrecht, 2012) (with one of the densest innervation by melanopsin cells; Brown *et al*., 2010; Beier *et al*., 2020), but also non-photic behaviours such as modulation of mood, sleep, spatial memory and food intake (Huang *et al*., 2019, 2021; Shi *et al*., 2019; Fernandez *et al*., 2020), and (3) the ventrolateral geniculate nucleus (VLG) associated with visuomotor and other non-image forming functions (Jeannerod & Putkonen, 1971; Legg & Cowey, 1977; Harrington, 1997). The VLG can be further divided into a brainstem input processing medial part (VLGm), and a directly retinorecipient lateral division (VLGl) (Niimi *et al*., 1963; Takatsuji & Tohyama, 1989; Kolmac & Mitrofanis, 2000).

Early studies show daily rhythms in the multi-unit LGN activity dependent on the SCN (Inouye & Kawamura, 1979) and an increased glucose utilisation in the LGN during the behaviourally active night (Jay *et al*., 1985; Room & Tielemans, 1989). This is supported by the daily variability in its spontaneous firing *in vivo* under urethane anaesthesia (Brown *et al*., 2011). Moreover, the IGL and VLG have been reported to stay in the reciprocal connection with the central clock (Watts & Swanson, 1987; Watts *et al*., 1987; Card & Moore, 1989; Moga & Moore, 1997; Moore *et al*., 2000). Accumulating evidence questions the exclusive role of the SCN as the circadian clock and demonstrates rhythmic clock gene expression in local brain clocks in the peripheral tissues (Granados-Fuentes *et al*., 2006; Guilding & Piggins, 2007; Herichová *et al*., 2007; Guilding *et al*., 2009, 2010; Zhang *et al*., 2014; Myung *et al*., 2019, 2018; Chrobok *et al*., 2020, 2021*c*; Northeast *et al*., 2020; Chrobok *et al*., 2021*d*, 2021*a*; Paul *et al*., 2020). However, despite a striking day-to-night difference in the main input to the LGN and a well-established involvement of the IGL in circadian processes (i.e., the SCN clock resetting), little has been known about its circadian physiology including intrinsic circadian timekeeping properties.

Here, we provide compelling evidence for intrinsic circadian timekeeping properties of the identified structures of the LGN. We study their clock genes expression *in vivo* as well as in isolated slice culture conditions. Additionally, we perform long-term multi-electrode array (MEA) recordings *ex vivo* combined with timed optogenetic manipulation of the retinal input to elucidate circadian variability in their neuronal firing and reveal the impact of photo-entrainment on their circadian activity pattern. Our study identifies novel circadian extra-SCN oscillators, with the DLG displaying clock gene expression exclusively *in vivo*, while the IGL and VLG possessing endogenous timekeeping mechanisms (seen also *ex vivo*). These are additionally reflected in the nocturnal elevation of their neuronal activity. Interestingly, we suggest that time of IGL peak firing is not shifted by retinal input in contrast to the VLG, which we propose as a new light-entrainable oscillator.

## MATERIALS AND METHODS

### Ethical approval

Experiments on rats were approved by the Local Ethics Committee in Krakow and animals were maintained and used according to Polish regulations and the European Communities Council Directive (86/609/EEC). Procedures carried out on mice were reviewed and approved by the Institutional Animal Care and Use Committee of Taipei Medical University (IACUC Approval No: LAC-2019-0118). All possible efforts were made to minimise the number of animals used and their sufferings.

### Animals

This study was performed on 94 adult, male Sprague Dawley rats. Animals were kept under standard 12:12 h light-dark (LD) cycle, unless stated otherwise, with *ad libitum* access to food and water. Rats were bred in the Animal Facility at the Institute of Zoology and Biomedical Research, Jagiellonian University in Krakow, housed three to six per cage at 23 ± 2°C and 67 ± 3% relative humidity. Four PERIOD2::LUCIFERASE (PER2::LUC) mice (RRID: IMSR_JAX:006852) were kept under a 12:12 light-dark cycle at 21 ± 2°C and 59 ± 4% relative humidity in Taipei Medical University Laboratory Animal Centre. All procedures in darkness were performed in infra-red night vision goggles (Pulsar, Vilnius, Lithuania).

### Quantitative reverse transcription PCR

#### Tissue preparation

Three separate cohorts of animals were subjected to the RT-qPCR study: (1) 24 rats were culled in four daily time points under LD cycle (n=6 each Zeitgeber time: ZT0, 6, 12 and 18), (2) 24 under DD at four circadian time points (n=6 each circadian time: CT0, 6, 12 and 18), and (3) 30 under DD at six circadian time points (n=5 for CT0, 4, 8, 12, 16 and 20). Two rats from the last group (one at CT8 and one at CT16) were excluded from analysis due to outlying results in all parameters measured. Animals culled in DD were moved to constant darkness conditions for two days before the cull. After establishing the deep anaesthetic state with isoflurane (2 ml/kg body weight; Baxter, USA), rats were decapitated and brains were quickly excised from the skull to ice-cold oxygenated (95% oxygen, 5% CO_2_) preparation artificial cerebro-spinal fluid (ACSF), composed of (in mM): NaHCO_3_ 25, KCl 3, Na_2_HPO_4_ 1.2, CaCl_2_ 2, MgCl_2_ 10, glucose 10, sucrose 125 and phenol red 0.01 mg/l. Brains were trimmed, mounted in the chamber of a vibroslicer (Leica VT1000S, Germany) and cut in 250 μm thick thalamic coronal sections. Bilateral LGNs were dissected using a scalpel and collected as whole, or cut immediately above the IGL to separate the DLG from the IGL+VLG. Subsequently, the tissue was flash frozen upon the dry ice and stored in −80°C, for up to a week. During the whole procedure, all instruments and surfaces were treated with RNaseZAP (Sigma, Germany) to block ribonuclease activity.

#### RNA isolation and RT-qPCR

Following the sampling, the RNA was extracted from the collected tissue with ReliaPrep RNA Tissue Miniprep System (Promega, USA). Obtained RNA was stored at −80°C in RNase-free water, before being processed to reverse-transcription using the High-Capacity RNA-to-cDNA Kit (Applied Biosystems, USA). The normalised amount of RNA was used for each sample. Subsequently, qPCR was carried out using PowerUp SYBR Green Master Mix (ThermoFisher Scientific, Lithuania) and StepOnePlus Real-Time PCR System (Applied Biosystems). For transcript amplification, QuantiTect primer assays (Qiagen, Germany) for selected genes were used. *Per1, Arntl* (*Bmal1*) and *Nr1d1* (*Reverbα*) were chosen to measure clock genes and *Gapdh* served as a housekeeping gene. Results were then analysed according to the Livak method (2^-ΔΔCT^) with *Gapdh* as reference gene and presented as relative target gene expression (RQ), where RQ=1 indicates the ZT/CT0 mean.

### PER2::LUC bioluminescence measurement

Four adult heterozygous PER2::LUC mice on a C57BL/6 background were deeply anesthetized under isoflurane and culled during the daytime (between ZT6 and ZT10). Brains were quickly isolated and transferred to ice-cold HBSS (ThermoFisher Scientific, MA) and prepared into 300 μm-thick coronal slices on a vibroslicer (Leica VT1000S, Germany). Slices were dissected into DLG and IGL+VLG under a stereomicroscope (Nikon, Japan) and transferred to culture membranes (Millipore Millicell-CM, MA). The explants were cultured in 35mm-dish (Dow Corning, MI) under the DMEM-based modified medium containing: phenol-red-free high-glucose DMEM (Sigma), 2% B-27 supplement (Gibco), 4.2 mM sodium bicarbonate (Gibco), 10 mM HEPES (Gibco), 1% penicillin-streptomycin (Gibco), and 300 μM beetle luciferin (Promega, WI). The dishes were sealed with silicone vacuum grease (Dow Corning, MI). PER2::LUC bioluminescence was continuously monitored with the LumiCycle (Actimetrics, IL). The data were exported and analyzed off-line with Mathematica (Wolfram Research, IL) using a custom-written package for spectral analysis and visualization (Myung *et al*., 2015).

### Viral vector injections

Seven 5-week old Sprague Dawley rats were deeply anaesthetised with isoflurane inhalation (3% v/v air mixture; Baxter, USA) in the flow chamber (Isoflurane Vaporizer; Stoelting, IL, USA). Following this, rats were given subcutaneous injections of Torbugesic (0.1 mg/kg body weight; Zoetis, USA) and Tolfedine 4% (4 mg/kg body weight; Biowet, Poland), while constantly exposed to isoflurane-air mixture through the breathing mask. Intraocular injections of AAV2-Syn-Chronos-GFP (2 μl, 2.1 × 10^12^ virus molecules/ml saline; UNC Vector Core, USA) were performed with 5 μl Hamilton syringe through 30G injection needle into a vitreous chamber of both eyes. Animals were returned to the animal facility and their condition was monitored. After four to five weeks post-injection, rats were subjected to further electrophysiological procedures.

### Electrophysiology

#### Tissue preparation

A total of ten rats were anaesthetised with isoflurane inhalation (2 ml/chamber) and culled at ZT0. Then, 250 µm thick acute thalamic coronal slices containing LGN were obtained in the same way as in *Quantitative Real-Time PCR, Tissue preparation* section. Slices were subsequently transferred to warm (32°C), carbogenated recording ACSF, composed of (in mM): NaCl 125, NaHCO_3_ 25, KCl 3, Na_2_HPO_4_ 1.2, CaCl_2_ 2, MgCl_2_ 2, glucose 5, phenol red 0.01 mg/l and penicillin-streptomycin 1 mg/ml (Sigma). Sections were allowed to rest for one hour before being transferred to the recording chamber of the multi-electrode array (MEA).

#### Recording

Spontaneous firing was evaluated with the use of the two-well perforated MEA *ex vivo* technology (Belle *et al*., 2021), by 30 h-long recordings of six LGN slices obtained from three 8-9 week old naïve rats and 14 slices from seven 9-10 week old rats injected with AAV2-Syn-Chronos-GFP. Slices were transferred to the recording wells of the MEA2100-System (Multichannel Systems GmbH, Germany) and positioned upon the 6×10 recording array of the perforated MEA (60pMEA100/30iR-Ti, Multichannel Systems). Sections were constantly perfused (2 ml/min) with fresh recording ACSF warmed to 25°C and gently sucked down the array by a vacuum pump connected to the underneath ASCF flow circuit throughout the whole experiment. Recordings were initiated at projected ZT (pZT) 2, after approximately half an hour of tissue settlement. Data were collected with Multi Channel Experimenter software (sampling frequency = 20 kHz; Multichannel Systems) repeatedly for 1 min in 10 min intervals.

#### Optogenetic stimulations

A blue diode (PlexBright LED 465 nm controlled by a LD1 LED driver; Plexon, Dallas, TX, USA) was coupled to an optical fibre (200 μm core diameter, 0.5 NA; Thorlabs, Bergkirchen, Germany). The end of the fibre was directed above the LGN slice in a distance enabling the beam of blue light to cover the whole recording area (Fig. 7B). The same light intensity (∼5 mW; measured with a photodiode power sensor S121C connected to a digital power meter PM100D; Thorlabs) and flash duration (1 ms) was used in all experiments. The stimulation protocol consisted of 15 trains of flashes (intra-train frequency: 10 Hz), applied in 2 min intervals. Thus, the whole stimulation protocol lasted 30 min. Custom-made scripts written in Spike2 (Cambridge Electronic Design Ltd, Cambridge, UK) were used to apply the optogenetic light stimulation protocol at pZT12.

#### Spike-sorting

Data were merged and then exported to HDF5 files with Multi Channel DataManager (Multichannel Systems GmbH). Subsequently, data were remapped and converted to DAT files with custom-made MatLab script (R2018a version, MathWorks). Finally, DAT files were automatically spike-sorted with KiloSort programme (Pachitariu *et al*., 2016) in MatLab environment. GPU (NVIDIA GeForce GTX 1050Ti GPU; CUDA 9.0 for Windows) was used for calculation speed improvement. In parallel, raw data were exported to CED-64 files (Spike2 8.11; Cambridge Electronic Design Ltd.) with Multi Channel DataManager, which were further filtered with Butterworth band pass filter (fourth order) from 0.3 to 7.5 kHz and remapped with a custom-made Spike2 script. Spike-sorting results were transferred into the prepared CED-64 files (using a custom-made MatLab script) for further visualisation and improvement. All spike-sorting results were manually revised in Spike2 8.11 with the aid of autocorrelation, principal component analysis and spike shape inspection to refine automatic sorting.

#### Analysis and data visualisation

To study circadian changes in firing rate, the single-unit activity (SUA) was 60 s binned and smoothed (Gaussian filter, width: 5 bins) in NeuroExplorer 6 (Nex Technologies, USA), which represented 10 mins of an actual recording. After that, the time of predominant peak activity was manually selected. These units which displayed several activity peaks, a dramatic fall of activity soon after the recording initiation or remained tonic throughout the recording, were excluded from peak analysis. Mean peak time (circular mean), peak clustering (Rayleigh test) and differences in peak distribution amongst studied areas (Watson-Williams test) were analysed and visualised with a Circular Statistics toolbox in MatLab.

Activity heatmaps and average firing plots were generated with custom-made MatLab script. For heatmaps, SUA was normalised for each unit separately, so the maximal firing = 1. Units were then sorted by the time of manually selected peaks (units without classified peaks were pulled down in a random order) or by the normalised activity during the projected night. Time of peak histograms were calculated in 1 h bins in Prism 7 (GraphPad Software, USA) and presented as relative frequency.

To assess the responsiveness of single units to optogenetic stimulation, peri-stimulus time histograms were prepared in NeuroExplorer 6. Neurons whose firing rate immediately following the light flash exceeded 95% confidence interval were deemed sensitive to the stimulation.

### Hybridisation *in situ*

#### Tissue preparation

Six adult Sprague Dawley rats were deeply anaesthetised with isoflurane and culled by decapitation (n=3 each Zeitgeber time: ZT6 and ZT18). Brains were then quickly removed from the skull, flash frozen upon the dry ice and stored at −80°C. They were subsequently cryo-sectioned at −20°C on a cryostat (Leica CM1950), thaw-mounted on Superfrost-Plus slides (Fisher Scientific, USA) and stored at −80°C overnight. On the next day, sections were fixed in cold (4°C) 4% paraformaldehyde (PFA) solution in phosphate-buffered saline (PBS) for 15 mins, rinsed in fresh PBS and dehydrated in increasing ethanol concentrations (50, 70, 100, 100%). Following, slides were let to air-dry and each slice was drawn around with a hydrophobic barrier pen.

#### RNAscope assay and imaging

Immediately following the tissue preparation, slices were processed with the RNAscope multiplex *in situ* hybridisation protocol (Advanced Cell Diagnostics – ACD, USA). First, slices were pre-treated with protease IV for 15 mins, rinsed in PBS and incubated for 2 h at 40°C with probes targeting *Per2* and *Penk*. Following, sections were rinsed with wash buffer and transferred to a four-step signal amplification protocol, terminating with the fluorophore tagging (*Per2* with Atto 550 and *Penk* with Atto 647). Finally, sections were rinsed twice with the wash buffer, stained with DAPI and coverslipped with the fluorescent mounting medium (ProLong™ Gold antifade reagent, Invitrogen, USA). Two LGNs per animal were imaged under 20 × magnification with the epifluorescence microscope (Axio Imager.M2, Zeiss, Germany) and images were inspected in ZEN software (ZEN 2.3. blue edition, Zeiss).

### Statistics

Statistical analysis of RT-qPCR results was performed in Prism 7. Daily and circadian variability was tested with ordinary one-way ANOVA or Kruskal-Wallis test, depending on the data distribution. MEA data were statistically analysed in MatLab using the Circular Statistics toolbox. Rayleigh test was chosen to assess the significance in peak clustering, whereas the Watson-Williams test (the circular analogue of ANOVA) was used to compare the peak distribution amongst the studied structures. Fisher’s test in Prism 7 was used to assess differences in proportion of cells responsive to optogenetic stimulation. In all tests, *p*<0.05 was deemed significant.

## RESULTS

### Rhythmic expression of core clock genes in the LGN in vivo

Limited evidence suggests daily variation in the LGN physiology (Paul *et al*., 2020), therefore we first aimed to establish if this thalamic complex of nuclei expresses core clock genes *in vivo*. We culled 24 rats housed in 12:12 LD conditions in four daily time points (with 6 h intervals) and measured the expression levels of *Per1* and *Bmal1* at ZT0, 6, 12 and 18. Notably, at the level of the whole LGN, there was a significant daily variability in the expression of these genes (*Per1*: *p*=0.0020, *Bmal1*: *p*=0.0392; Fig. 1A), with the *Per1* acrophase at the beginning of the night, coinciding with the nadir of *Bmal1* expression. Following this, we moved another cohort of 24 rats into constant darkness conditions (DD), ∼48 h before cull in four circadian time points (CT0, 6, 12 and 18), mimicking these daily ones. Again, we noticed a significant circadian variability in *Per1* (*p*=0.0004) and *Bmal1* (*p*=0.0024, Kruskal-Wallis tests; Fig. 1B) expression, with unchanged temporal patterning compared to LD condition. These observations suggest that the LGN expresses core clock genes in the rhythmic fashion independently of the light-dark cycle.

**Figure 1.**
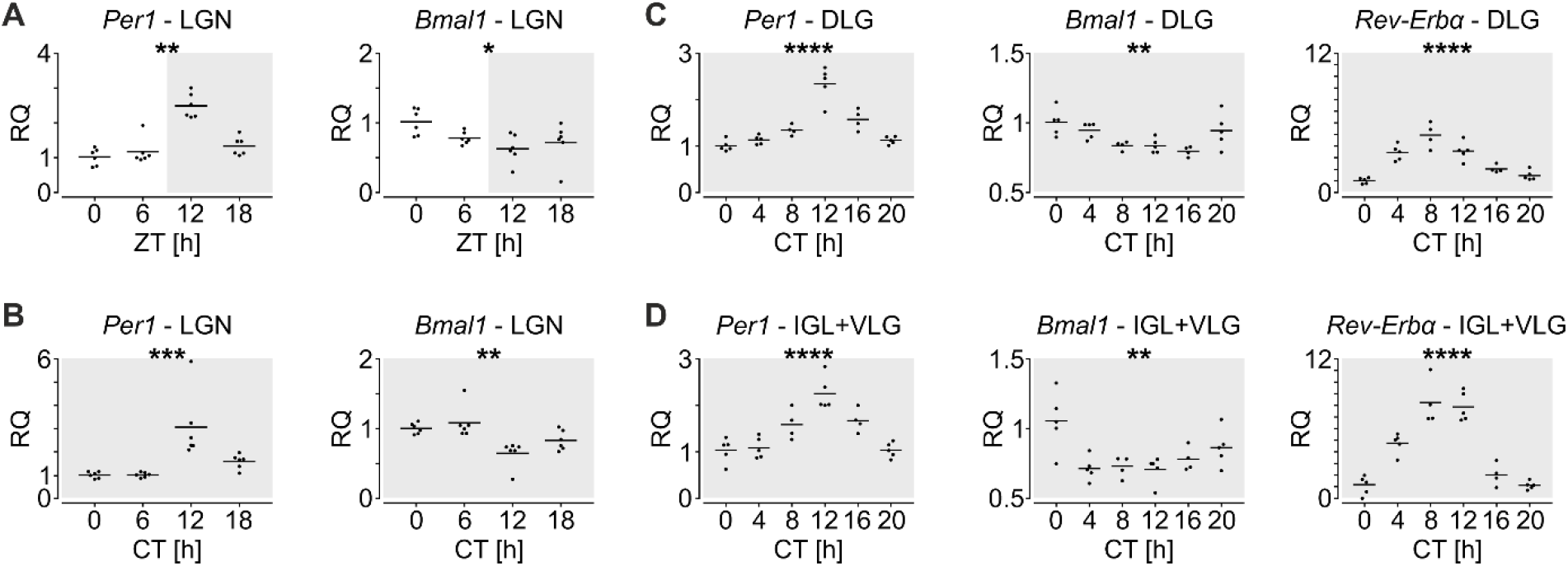
Rhythmic core clock gene expression in the lateral geniculate nucleus (LGN) *in vivo*. (**A**) Variation in *Per1* and *Bmal1* in the whole LGN in four daily time points of the 12:12 light-dark (LD) cycle. \*\**p*=0.0020, \**p*=0.0392, n=24 rats, Kruskal-Wallis tests. (**B**) Circadian variation in core clock gene expression in the whole LGN under constant darkness (DD). \*\*\**p*=0.0004, \*\**p*=0.0024, n=24 rats, Kruskal-Wallis tests. (**C**) Circadian variation in *Per1, Bmal1* and *Rev-Erbα* in the isolated dorsolateral geniculate nucleus (DLG) under DD, measured in six circadian time points. \*\*\*\**p*<0.0001, \*\**p*=0.0025, n=28 rats, one-way ANOVAs. (**D**) Circadian variation in clock genes in the isolated intergeniculate leaflet (IGL) together with the ventrolateral geniculate nucleus (VLG). \*\*\*\**p*<0.0001, \*\**p*=0.0020, n=28 rats, one-way ANOVAs. ZT – Zeitgeber time, CT – circadian time. Dark boxes code the dark phase.

To further elucidate if all or only a distinct subset of the LGN expresses clock genes *in vivo*, we repeated the RT-qPCR experiment with a higher spatio-temporal resolution. Therefore, 30 rats were moved to DD for two days before being culled in six circadian time points (CT0, 4, 8, 12, 16 and 20). Subsequently, we dissected and analysed the LGN in two separate samples – one containing the DLG and the other the IGL together with the VLG. In both the DLG and IGL+VLG, we found *Per1* and *Bmal1* to exhibit circadian variability (*Per1*: *p*<0.0001, *Bmal1*_*DLG*_: *p*=0.0025, *Bmal1*_*IGL+VLG*_: *p*=0.0020; Fig. 1C,D) in the relation that approximates antiphase. Additionally, we measured circadian transcript levels of *Reverbα*, a key regulating factor for *Bmal1* expression, forming a secondary feedback loop (Ikeda *et al*., 2019). Surprisingly, *Reverbα* expression displayed a robust circadian change in both studied regions of the LGN (*p*<0.0001, one-way ANOVAs; Fig. 1C,D), five to eight-fold higher in its peak at CT8 compared to a trough at CT0. The maximal expression of *Reverbα* preceded the *Per1* peak by four hours. In the IGL+VLG it stayed relatively high at CT12, whereas in the DLG the *Reverbα* expression gradually declined throughout the night. Altogether this dataset provides evidence for rhythmic clock gene expression in both dorsal and ventral parts of the LGN *in vivo*.

### Intrinsic circadian timekeeping in the LGN ex vivo

The mismatch between the expression of clock genes *in vivo* and the intrinsic activity to sustain their rhythmic expression *ex vivo* has been reported before (Yoo *et al*., 2004; Mei *et al*., 2018). Therefore, to exclude this possibility, we next cultured brain slices obtained from PER2::LUC mice (n=4) containing exclusively the DLG or the IGL together with VLG, and recorded bioluminescence for one week. Explants comprising the DLG expressed very weak or negligible circadian fluctuation in PER2::LUC expression (Fig. 2A). However, those containing IGL+VLG displayed notable circadian rhythmicity in bioluminescence throughout the whole recording duration, with the period close to 24 h (Fig. 2B). These results indicate an intrinsic nature of clock gene expression in the IGL+VLG, but not the DLG.

**Figure 2.**
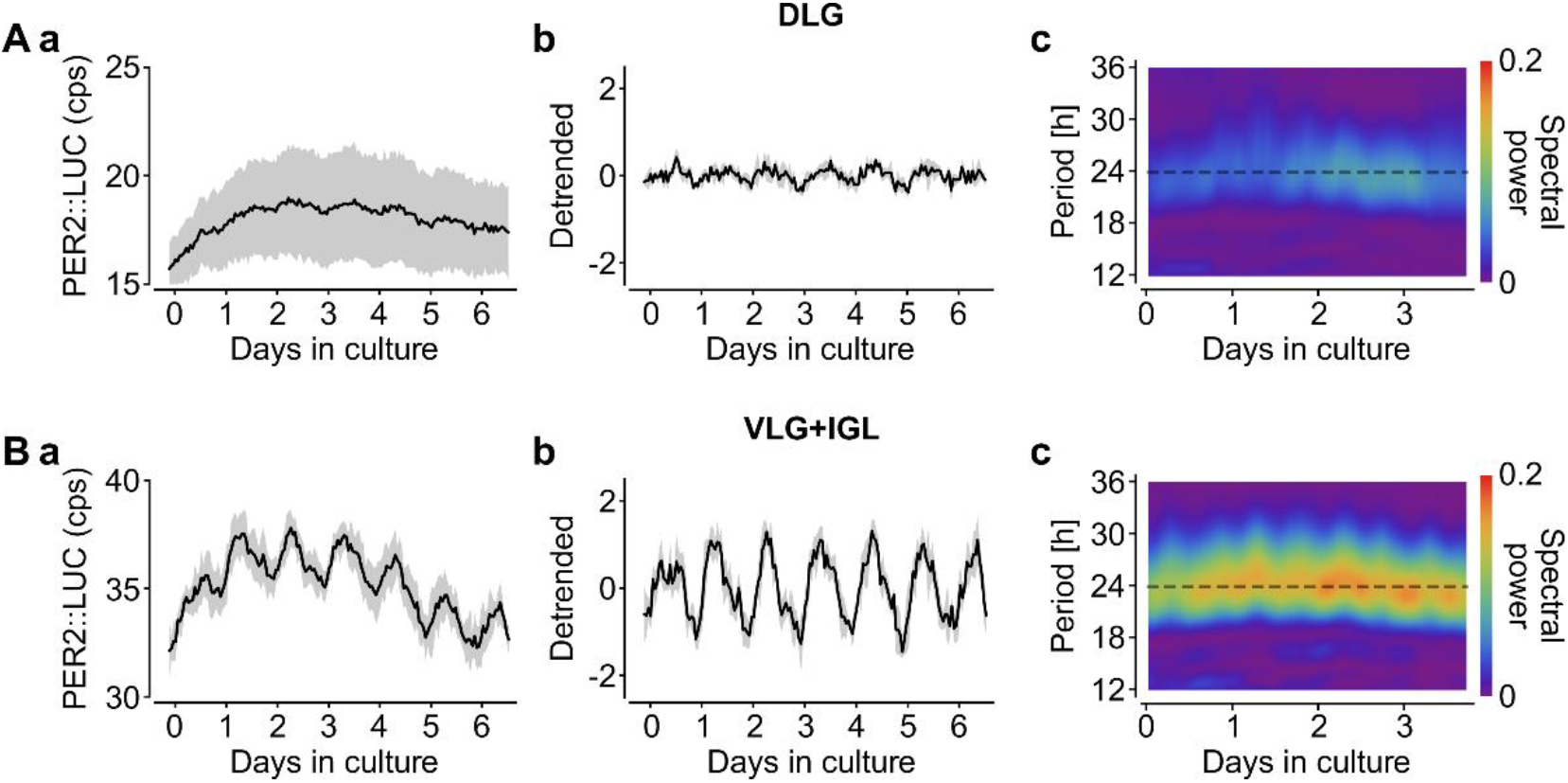
Intrinsic PER2::LUC bioluminescence oscillations in the ventrolateral geniculate nucleus (VLG) and intergeniculate leaflet (IGL) in slice culture. (**A**) PER2::LUC expression in the dorsolateral geniculate nucleus (DLG). (**B**) PER2::LUC bioluminescence in the VLG+IGL. (**a**) Raw bioluminescence signal. The solid line represents an ensemble average (n=4 animals) and the shade indicate the standard deviation of the ensemble mean. (**b**) Detrended PER2::LUC signal. (**c**) The spectrogram through the sliding fast Fourier transformation (FFT) reveals a circadian spectral power around 24-h period (ensemble average over spectrograms, n=4 animals). Note a strong spectral power in the VLG+IGL and a weak one in the DLG.

### The IGL and VLG neurons elevate their neuronal firing during the projected night

To communicate the circadian phase with other neurons within and between brain areas, molecular circadian clocks in the SCN and extra-SCN oscillators are accompanied by daily changes in their neuronal activity (Granados-Fuentes *et al*., 2004; Belle *et al*., 2009; Guilding *et al*., 2009; Sakhi *et al*., 2014; Chrobok *et al*., 2020, 2021*d*, 2021*c*, 2021*a*, 2021*e*). To assess if the LGN firing rates are similarly coordinated across the circadian cycle, we next performed ∼30 h-long multi-electrode array (MEA) recordings of neuronal activity from six thalamic brain slices. The LGN was placed over the recording electrodes to encompass the DLG, IGL and VLG, which could be further divided into brainstem information processing medial part (VLGm) and retinorecipient lateral division (VLGl). No evident circadian change in the single-unit activity (SUA) was observed in the DLG (Fig. 3A), which paralleled the lack of intrinsic PER2::LUC rhythmicity in this area. On the contrary, the mean SUA in the IGL displayed a prominent rise beginning at late projected day and peaking at around pZT17-18 (Fig. 3B). Similarly, the averaged neuronal activity in the VLG started to elevate at early projected night, reaching the peak at pZT18-19 in the VLGm (Fig. 3C) and pZT20 in the VLGl (Fig. 3D). This spatio-temporal pattern was also observed in the multi-unit activity of the LGN subnuclei (Fig. 4; SMovie 1,2).

**Figure 3.**
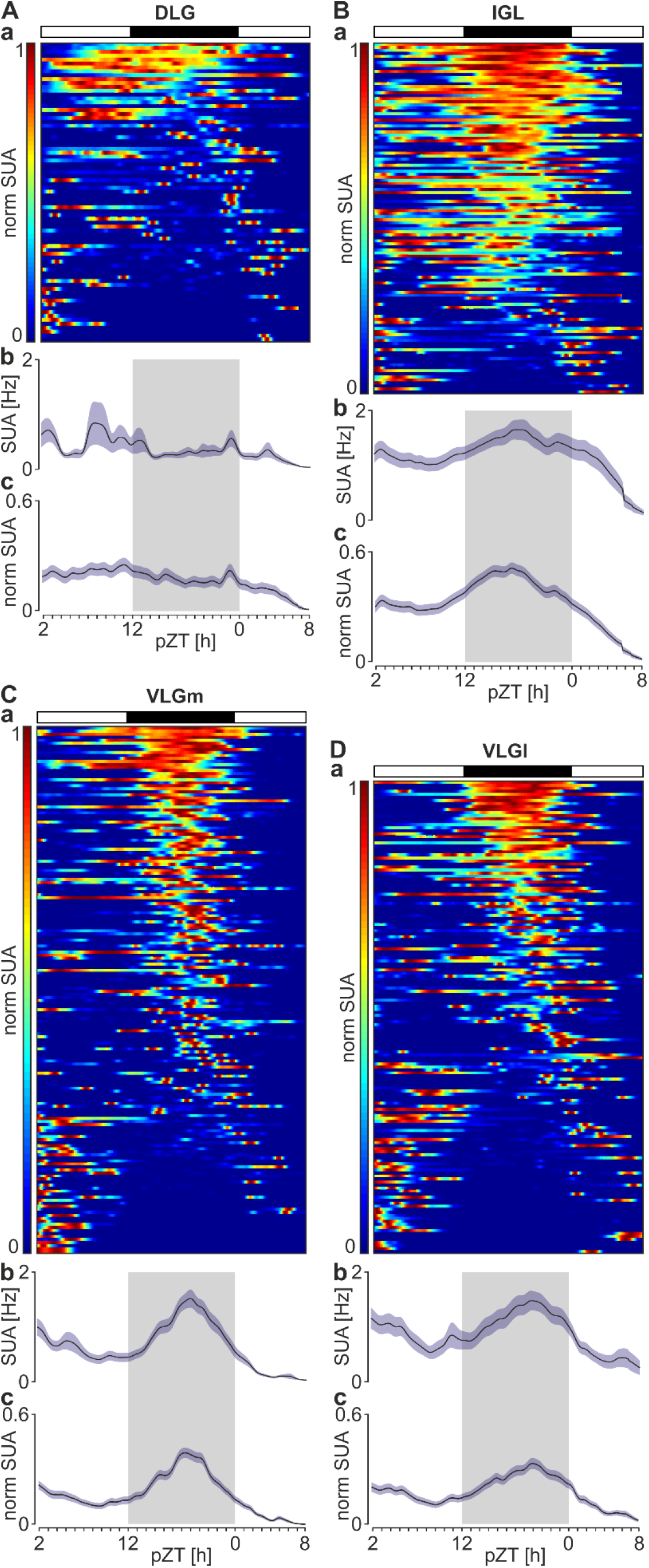
The intergeniculate leaflet (IGL) and ventrolateral geniculate nucleus (VLG) display an increase in neuronal activity during the projected night *ex vivo*. Panels collect results for the dorsolateral geniculate nucleus (DLG, ***A***), IGL (***B***), and two parts of the VLG: medial (VLGm, ***C***) and lateral (VLGl, ***D***). (**a**) Heatmaps showing the normalised single unit activity (SUA) for each recorded neuron (bin: 10 min) over the course of 30 h recording on the multi-electrode arrays *ex vivo*, segregated from the most (*top*) to the least active during the night (*bottom*). Warm colours code the relatively high, whereas cold – low firing rates. (**b**) Average plot (bin: 10 min) of the raw SUA. (**c**) Average plot (bin: 10 min) of the normalised SUA. Data were drawn as mean ± SEM. Note the nocturnal rise of neuronal activity in the IGL and VLG, but not the DLG. pZT – projected Zeitgeber time. n=523 units from six slices obtained from three rats.

**Figure 4.**
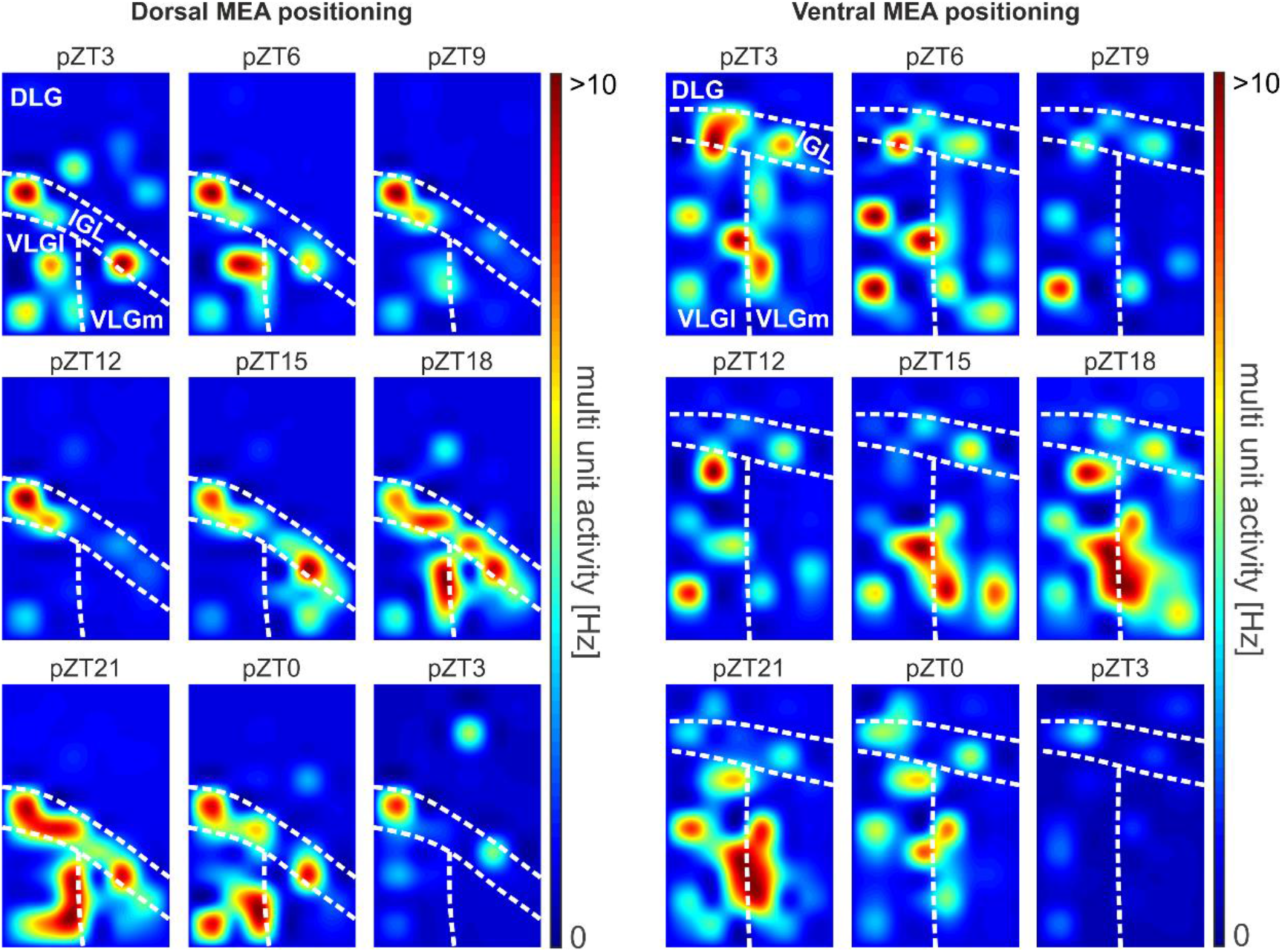
Spatio-temporal distribution of multi-unit neuronal activity in the lateral geniculate nucleus (LGN). Heatmaps demonstrating multi-unit firing rates from each recording location in the dorsolateral geniculate nucleus (DLG), intergeniculate leaflet (IGL) and two parts of the VLG: lateral (VLGl) and medial (VLGm). Warm colours indicate high firing rates, whereas cold colours indicate neuronal silence. The dotted line delineates the borders of LGN substructures. pZT – projected Zeitgeber time.

Subsequently, we manually assessed the time of the highest firing for each single unit recorded, omitting those displayed a fast drop in SUA after the initiation of the recording, showed multiple peaks (mostly in the DLG) or remained tonic throughout the experiment (most common for the IGL) (Fig. 5A-D). Circular analysis revealed that SUA peaks were not significantly grouped around any particular circadian time for the DLG (*p*=0.6154), but showed a prominent clustering for the IGL and both parts of the VLG (*p*<0.0001, Rayleigh tests; Fig. 5E). In detail, the IGL SUA peaked at pZT18.3, followed by the VLGm at pZT19, and finally by VLGl at pZT21.2 (data presented as circular means). The difference in peak SUA amongst the rhythmic parts of the LGN was additionally validated by the Watson-Williams test (the circular equivalent of ANOVA) indicating a significant variability between the IGL and VLGl (*p*=0.0005) and between two parts of the VLG (*p*=0.0017; Fig. 5E). Taken together, the results of our electrophysiological experiments provide another layer of evidence on the intrinsic ability of IGL and VLG neurons to convey timekeeping information provided by rhythmic clock gene expression through the circadian variation in the neuronal activity.

**Figure 5.**
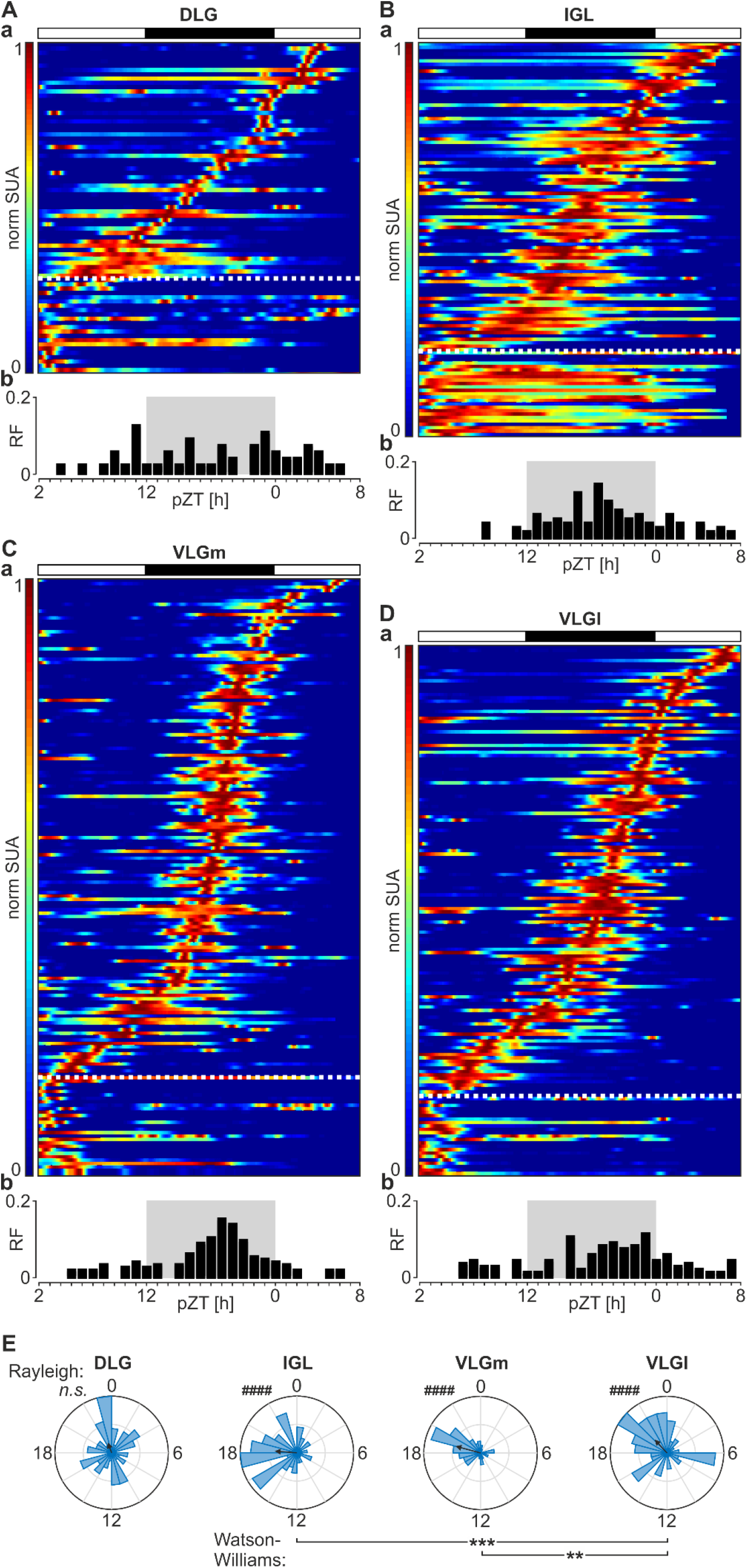
Temporal distribution of peak activity in the lateral geniculate nucleus (LGN). Panels encapsulate results for the dorsolateral geniculate nucleus (DLG, ***A***), IGL (***B***), and two parts of the VLG: medial (VLGm, ***C***) and lateral (VLGl, ***D***). (**a**) Heatmaps showing the normalised single unit activity (SUA) for each recorded neuron (bin: 10 min) over the course of 30 h recording on the multi-electrode arrays *ex vivo*, segregated by the time of a manually selected predominant peaks in activity, from the latest to the earliest. Warm colours code the relatively high, whereas cold colours code low firing rates. Non-rhythmic units are separated by the dotted horizontal line and pulled below. (**b**) Histograms of peaks (bin: 1 h). (**E**) Rose petal diagrams (circular histograms of peaks) for each substructure (bin: 1 h). The length of the vector codes the strength of peak clustering. n.s. *p*=0.6154, ####*p*<0.0001, Rayleigh tests. \*\*\**p*=0.0005, \*\**p*=0.0017, Watson-Williams tests (circular analogue of ANOVA). pZT – projected Zeitgeber time. n=523 units from six slices obtained from three rats.

### Clock gene Per2 is expressed throughout the LGN but not by enkephalinergic neurons

A clear distinction between the parts of the LGN is assured by the IGL, which intercalates between the DLG and VLG (Moore & Card, 1994). It is populated by retinorecipient enkephalinergic neurons forming the geniculo-geniculate pathway and neuropeptide Y (NPY)-synthesising cells, which do not receive retinal innervation yet send their axons to the SCN. Whilst the NPY neurons are limited to the IGL, scattered enkephalinergic cells can be also found throughout the VLG, but not DLG (Harrington, 1997; Morin & Allen, 2006). Thus, we next aimed to investigate the pattern of clock gene expression in the LGN, focusing on its retinorecipient subpopulations. The fluorescent *in situ* hybridisation was performed with the RNAscope technology on LGN slices obtained from six rats, three culled in the middle of the day, and three in the middle of the night. Signals from fluorescent probes against *Per2* were found in all three parts of the LGN (Fig. 6). Notably, clusters of *Per2* in the LGN were not observed around each DAPI-stained nucleus, but rather the clock gene expressing cells were scattered across the structure. Further visual inspection of all slices revealed a negligible, if any, co-localisation of *Per2* with *Penk* (the transcript for proenkephalin), neither in the IGL (Fig. 6b,c) nor VLG (Fig. 6d,e). These results suggest that rhythmic clock gene expression in the IGL and VLG may be carried out by other, non-retinorecipient neuronal subpopulations.

**Figure 6.**
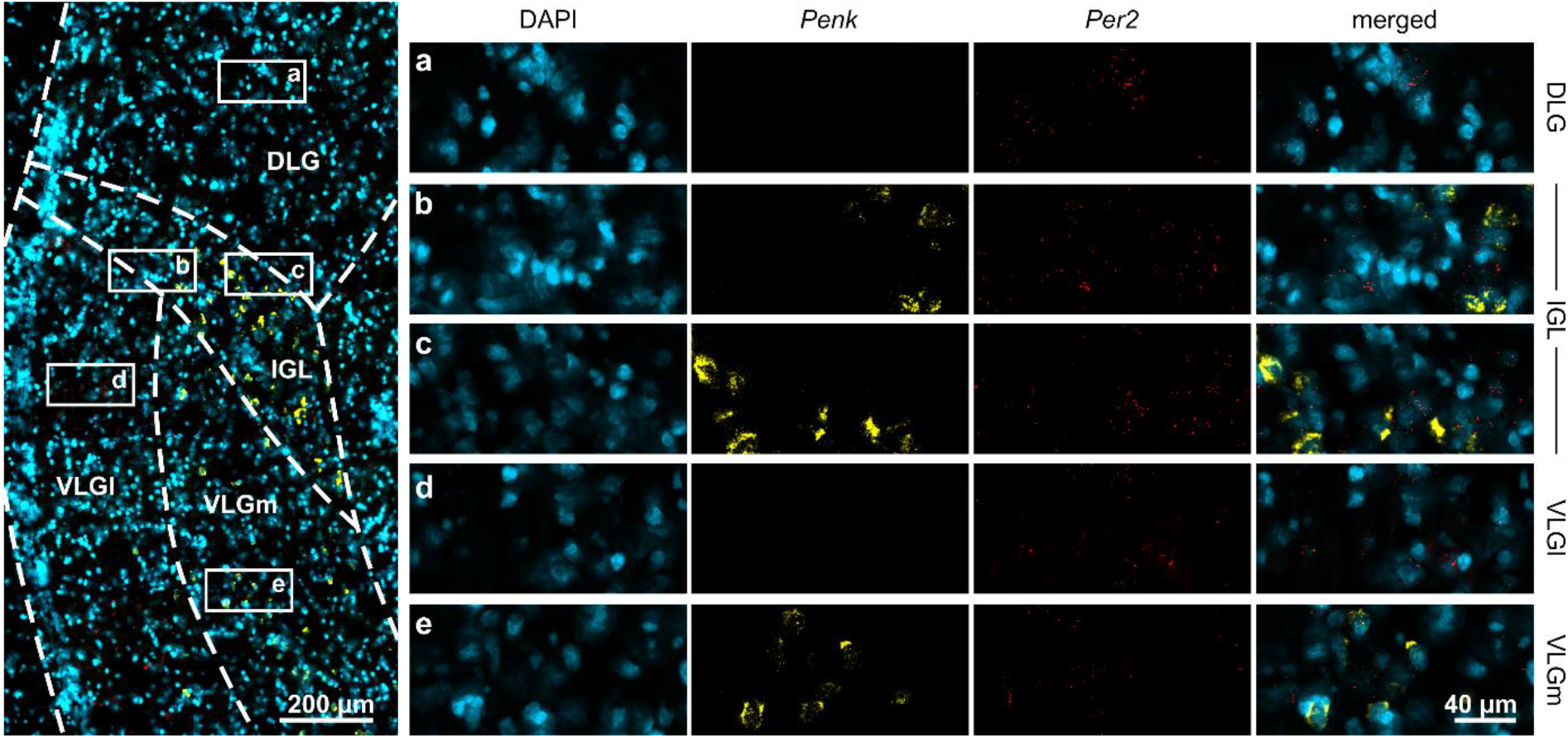
*Per2* expression in the lateral geniculate nucleus (LGN) by non-enkephalinergic cells. (***left panel***) Low magnification photomicrography with the LGN subnuclei outlined: the dorsolateral geniculate nucleus (DLG), intergeniculate leaflet (IGL), and two parts of the ventrolateral geniculate nucleus (VLG): and lateral (VLGl) and medial. (***right panel***) High magnification of inserts from: the DLG (***a***), IGL (***b***,***c***), VLGl (***d***) and VLGm (***e***). Nuclei counterstained with DAPI are presented in cyan, *Penk* (transcript for proenkephalin) in yellow and *Per2* in red. The merged signal shows no co-localisation of *Penk* and *Per2*.

### Circadian variability in neuronal activity of the VLG, but not IGL, is light-entrainable ex vivo

Clock cells in the SCN receive dense retinal innervation, which can phase-shift their molecular and electrical activity rhythms. Due to this feature, the term ‘light-entrainable oscillator’ has been coined for the central clock (Hastings *et al*., 2018, 2019). We next aimed to evaluate if circadian variability in the LGN neuronal activity can be entrained by the retinal input. Seven rats were intraocularly injected with AAV2-Syn-Chronos-GFP to both eyes, and after four to five weeks post-injection animals were culled for *ex vivo* MEA recordings combined with optogenetic stimulation of retinal terminals in the LGN.

First, optogenetic stimulation protocol was tested on two LGN slices (Fig. 7A,B) in an acute setup. The protocol consisted of 15 trains of flashes (1 ms long, 10 Hz), 1 minute duration each, separated by 1 minute silence. Therefore, the whole stimulation lasted 30 minutes. This stimulation pattern was designed to mimic the natural arrival of retinal excitatory post-synaptic currents seen *in vivo* where the retino-thalamic pathway is intact and the LGN activity follows the retinal infra-slow oscillation, with a period of about two minutes (Lewandowski & Błasiak, 2004; Blasiak & Lewandowski, 2013; Chrobok *et al*., 2018; Orlowska-Feuer *et al*., 2021). High frequency (beta/gamma) oscillations are also imposed by the retina on the LGN activity *in vivo* (Saleem *et al*., 2017; Storchi *et al*., 2017; Chrobok *et al*., 2018; Orlowska-Feuer *et al*., 2021); here we used lower frequency train stimulation (10 Hz), due to its highest fidelity in the slice preparation (Chrobok *et al*., 2021*b*). Indeed, our optogenetic protocol was efficient to excite LGN neurons throughout the whole 30 min-long stimulation (Fig. 7C).

**Figure 7.**
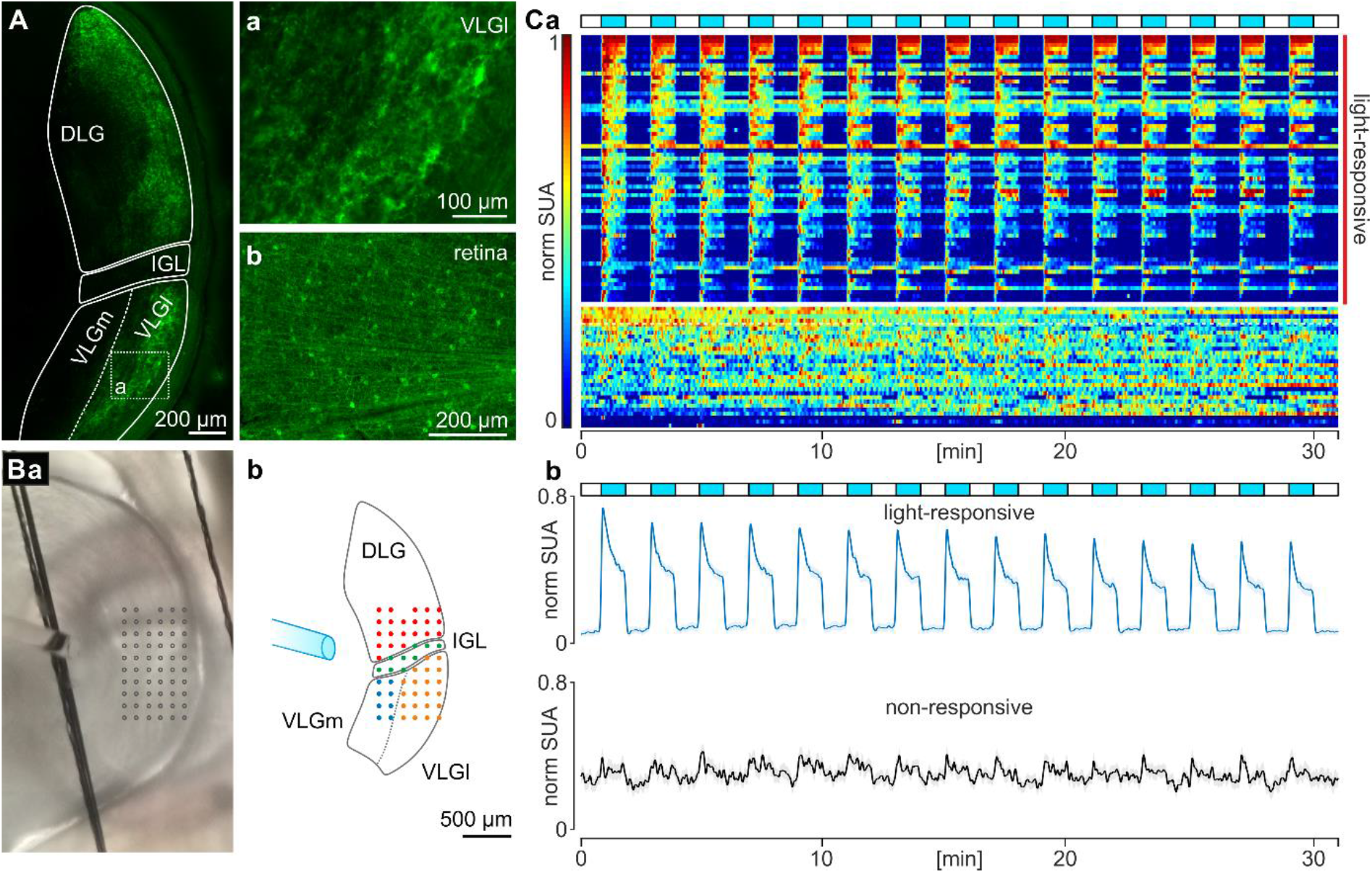
Optogenetic stimulation of retinal terminals in the lateral geniculate nucleus (LGN) *ex vivo*. (**A**) Example epifluorescence photomicrograph of the LGN slice after an intraocular injection of AAV-Syn-Chronos-GFP. (**Aa**) High magnification of axonal terminals at the area of ventrolateral geniculate nucleus, lateral division (VLGl). (**Ab**) Retinal preparation with GFP-positive cells and axonal projections. (**B**) Positioning of the brain slice containing LGN upon the 6 × 10 multi-electrode array (MEA), showing the end of the optic fibre targeting the recorded brain area. (**Ba**) Photograph with the MEA location indicated by grey circles. (**Bb**) Reconstruction with the borders of LGN substructures outlined. Red circles represent recording locations in the dorsolateral geniculate nucleus (DLG), green – in the intergeniculate leaflet (IGL), blue – in the medial division of the VLG, and orange – in the VLGl. (**C**) Optogenetic stimulation protocol test. (**Ca**) Heatmap displaying normalised single unit activity (SUA) throughout the stimulation protocol (bin: 1 s). Blue boxes code the time of the optogenetic stimulation (flash duration: 1 ms, intra-train flash frequency: 10 Hz), repeated 15 times in 2 min intervals. Units were sorted from top to bottom according to relative activity during the first response to optogenetic stimulation. Non-responsive units were pulled down and separated by a solid white horizontal line. (**Cb**) Average plots showing normalised SUA of all light-responsive (in blue) and non-responsive units (in black). Data were presented as mean ± SEM. n=97 units from two slices.

Next, we performed 30 h-long MEA recordings of twelve LGN slices obtained from AAV2-Syn-Chronos-GFP injected rats, initiated at pZT2. Six slices were stimulated at pZT12 with the optogenetic protocol described above, simulating the light pulse at the beginning of dark phase. Remaining six served as control and were not stimulated at any time. Consistent with the previous experimental set, the DLG firing rates did not display notable circadian variability at the whole area level, neither in control nor stimulated slices (Fig. 8A). This was attributable to a lack of temporal organisation in the time of single unit circadian activity peaks in this structure (control: *p*=0.1934, stim: *p*=0.8885, Rayleigh tests; Fig. 8Ea). Neurons localised in the IGL of the control slices exhibited an organised elevation in their firing rates, peaking at pZT14.9 (*p*<0.0001; Rayleigh tests; Fig. 8B,Eb). Optogenetic stimulation of retinal afferents to the LGN did not significantly change the time of the IGL peak (at pZT16.2, *p*=0.2664, Watson-Williams test; Fig. 8B,Eb), with the similarly organised peak time amongst its single units (*p*<0.0001, Rayleigh test; Fig. 8Eb). In contrast, the stimulation protocol at pZT12 was efficient in shifting the peak firing in both divisions of the VLG (Fig. 8C,D). In the densely retinorecipient VLGl, the mean neuronal activity peak was significantly time delayed by 2.9 h (control: pZT14.1, stim: pZT17, *p*<0.0001, Watson-Williams test; Fig. 8Ec), with no differences in peak clustering (*p*<0.0001, Rayleigh tests, Fig. 8Ec). The VLGm also exhibited a robust 4.5 h phase delay in its peak neuronal activity upon the stimulation of retinal terminals, with a mean peak at pZT11.9 in control conditions *vs* pZT16.4 in stimulated slices (*p*<0.0001, Watson-Williams test; Fig. 8Ed). Control VLGm cells exhibited a clustered peak distribution (*p*<0.0001), whereas the optogenetic stimulation tended to decrease their synchrony (*p*=0.0115, Rayleigh tests, Fig. 8Ed), suggesting only a distinct subpopulation of neurons shift their firing phase. Altogether, our observations suggest the VLG to retain light-entrainable oscillators, in contrast to the IGL, whose circadian phase remains unchanged after the activation of retinal input.

**Figure 8.**
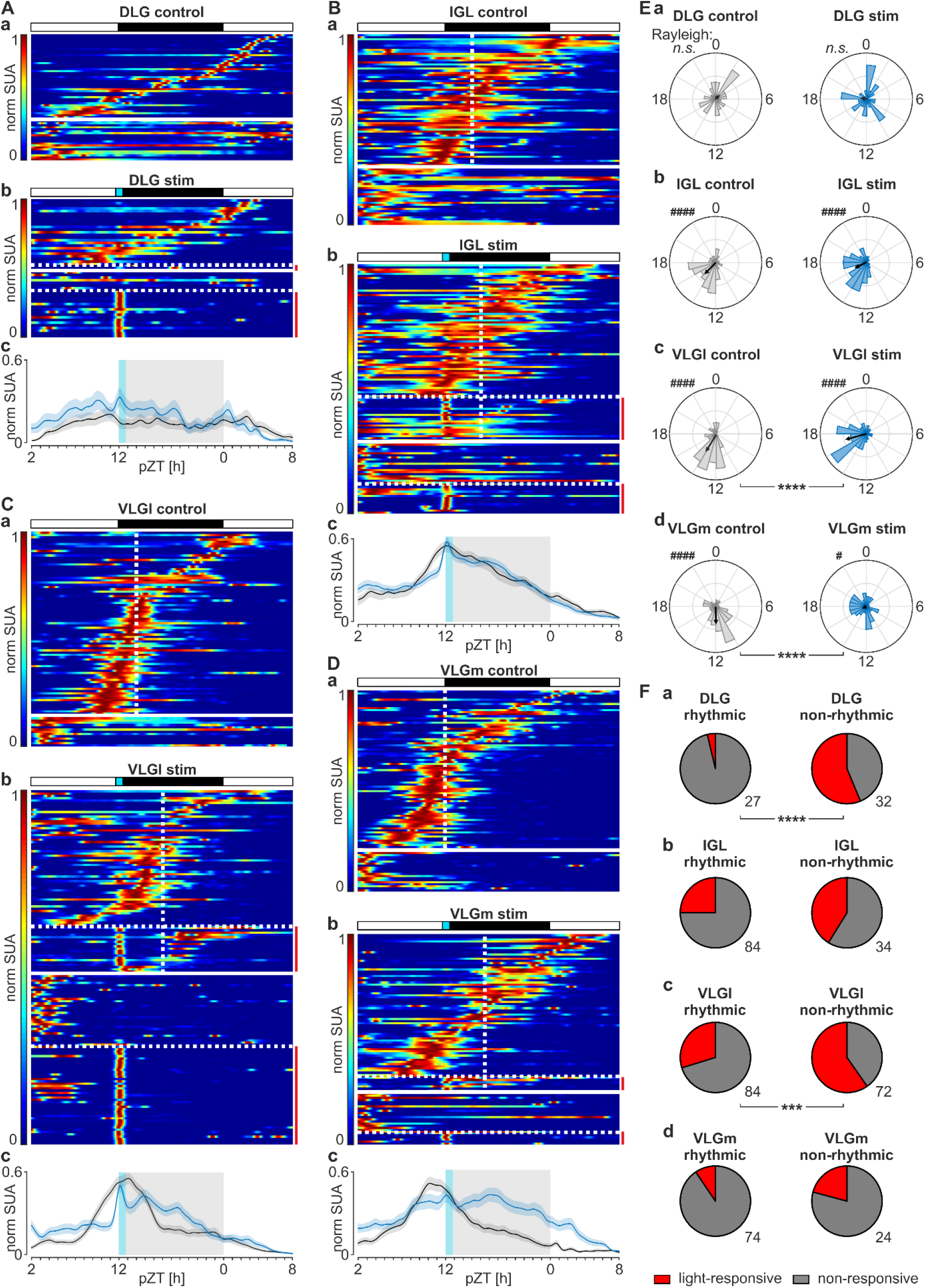
Circadian variation in neuronal activity is light-entrainable in the ventrolateral geniculate nucleus (VLG), but not the intergeniculate leaflet (IGL). Panels collect results for the dorsolateral geniculate nucleus (DLG, ***A***), IGL (***B***), and two parts of the VLG: lateral (VLGl, ***C***) and medial (VLGm, ***D***). (**a**,**b**) Heatmaps showing the normalised single unit activity (SUA) for each recorded neuron (bin: 10 min) over the course of 30 h recording on the multi-electrode arrays *ex vivo*, segregated by the time of a manually selected predominant peaks in activity, from the latest to the earliest. Warm colours code the relatively high, whereas cold – low firing rates. Subpanel ***a*** demonstrates results from control slices, while ***b*** – from optogenetically stimulated ones. Non-rhythmic units are separated by the solid horizontal line and pulled below. Units responsive to optogenetic stimulation were separated by dotted horizontal lines and pulled below in both rhythmic and non-rhythmic group. Vertical dotted line points the mean time of peak. Blue rectangle shows the time of a 30 min-long optogenetic stimulation (described in Figure 7) at projected Zeitgeber time (pZT) 12. (**c**) Average plots displaying normalised SUA for the control (in black) and stimulated slices (in blue). Data were presented as mean ± SEM. (**E**) Rose petal diagrams (circular histograms of peaks) for rhythmic units for each substructure in control *vs* stimulated condition (bin: 1 h). The length of the vector codes the strength of peak clustering. DLG_control_ n.s. *p*=0. 1934, DLG_stim_ n.s. *p*=0. 8885, \**p*=0.0115, ####*p*<0.0001, Rayleigh tests. \*\*\*\**p*<0.0001, Watson-Williams tests. (**F**) Pie charts representing the proportion of light-responsive units (in red) to these non-responsive to optogenetic stimulation (in grey), for the rhythmic and non-rhythmic neurons in each LGN substructure. \*\*\**p*=0.0002, \*\*\*\**p*<0.0001, Fisher’s tests. Numbers in the bottom right present the number of cells recorded in each group from seven stimulated and seven control slices (n=7 rats).

Surprisingly, the minority of ‘rhythmic’ LGN cells (i.e., those exhibiting a single circadian activity peak) were sensitive to the optogenetic stimulation of retinal afferents. This proportion was higher in the subpopulation of ‘non-rhythmic’ neurons, which was significant for the DLG (*p*<0.0001) and VLGl (*p*=0.0002, Fisher’s tests; Fig. 8Fa,c), but not the IGL (*p*=0.1183) nor the sparsely light-responsive VLGm (*p*=0.1599, Fisher’s tests; Fig. 8Fb,d). These results suggest that directly retinorecipient cells transmit the information of photic entrainment upon the secondary subpopulation of LGN neurons that possess intrinsic circadian timekeeping properties.

## DISCUSSION

Our study reveals the thalamic LGN as a multicomponent circadian oscillatory centre with its distinct parts exhibiting different timekeeping properties. Here we propose the DLG to express organised clock gene rhythms exclusively *in vivo*, most probably imposed by other brain clocks, as they disappear in *ex vivo* conditions. More importantly, we report the IGL and VLG to sustain rhythmic clock gene expression in isolated conditions and to display an endogenous nocturnal rise in their neuronal activity. Additionally, we propose the VLG to act as a light-entrainable oscillator, in contrast to the IGL whose circadian firing rates are not phase shifted by the excitation from the retinal input.

Historically, the SCN was considered the only brain area to sustain rhythmic expression of clock genes (Hastings *et al*., 2018, 2019). However, with the breakthrough application of the luciferase reporter system, especially the construction of the knock-in PER2::LUC mouse model (Yoo *et al*., 2004), it has become clear that several brain areas and peripheral tissues display organised autonomous and semi-autonomous rhythms of core clock gene expression that last for several days, or even weeks in culture (Guilding & Piggins, 2007; Guilding *et al*., 2009; Paul *et al*., 2020; Begemann *et al*., 2020).

Recently, we discovered another extra-SCN retinorecipient centre – the superior colliculus (SC) – to rhythmically express *Per1, Bmal1* and *Reverbα* (Chrobok *et al*., 2021*a*). Here we show that both the DLG and the IGL+VLG rhythmically express core clock genes *in vivo*, with a characteristic phase relation seen in the SCN and other circadian oscillators (Takahashi, 2017). Rhythmic expression of clock genes was preserved in constant darkness conditions, which proves the independence of LGN clock mechanisms from the cyclic ambient light changes. Altogether, it suggests that clock mechanisms are at work in the whole LGN, leastwise *in vivo*, when the LGN is connected to its afferents. Notably, the *Per1* expression peaks at the light-dark transition (ZT12) or at the beginning of active phase (CT12), similarly to other extra-SCN oscillators such as the SC, circumventricular organs, or the dorsal vagal complex (Kaneko *et al*., 2009; Chrobok *et al*., 2020, 2021*a*; Northeast *et al*., 2020). Thus, this molecular rhythm is phase shifted from the central clock which together with oscillators like habenula peaks around the middle of the day (Guilding *et al*., 2010; Sakhi *et al*., 2014; Zhao *et al*., 2015; Takahashi, 2017; Baño-Otálora & Piggins, 2017). In the SCN, the phase gap between its dorsal and ventral part encodes the length of the light phase, therefore provides a seasonal calendar rather than a daily clock (Coomans *et al*., 2015; Myung *et al*., 2015; Myung & Pauls, 2018). However, the information coded by the phase difference amongst a network of brain oscillators is still elusive. The LGN network exhibits heterogeneous spatiotemporal dynamics under optogenetic emulation of retinal light input. The segregated organisation of retinorecipient input, autonomous clock and SCN efferent components suggests a sophisticated compartmentalisation of the network, which might be necessary since the LGN makes much more extensive connections compared to the SCN.

The essential question in the experimental chronobiology on how the molecular clock communicates with the electrical one at the level of neuronal membrane remains mostly unresolved (Allen *et al*., 2017; Belle & Allen, 2018; Belle & Diekman, 2018). It is essential for clock cells to communicate their phase in order to synchronise and generate a coordinated output. We showed that the IGL and both parts of the VLG are capable to produce circadian changes in the firing rate, even in isolated conditions of the *ex vivo* MEA recording. Long-term recordings used in this study create an advantage over the acute set-up, where neuronal activity is sampled across 24 h from animals culled at distinct daily time points. This is due to (1) an absence of photic and non-photic cues during the long-term recording, compared to behavioural challenges proceeding the cull in shorter-term set-up, and (2) potential clock resetting by the cull itself (Guilding *et al*., 2009). In our recent study, we used four daily time points to sample LGN firing rates across the daily cycle and found the VLG to elevate its firing at ZT18, although nocturnal elevation of neuronal activity was not significant for the IGL (Chrobok *et al*., 2021*b*). We hypothesise that the IGL may be more prone to phase resetting by non-photic cues such as acute arousal preceding the cull, compared to the VLG, or alternatively, its rhythmicity was not robust enough to appear in the short-term recording set-up.

We demonstrated the neuronal activity of the IGL and VLG to peak in the middle of the projected night – a time delayed from the peak *Per1* expression by approximately 6 h. In the SCN clock, the highest firing rates are observed in the middle of a day, temporarily aligned with peak *Per1* expression (Takahashi, 2017). It has been hypothesised that clock gene expression may be coupled to the activity of distinct ion channels, including a reduction of potassium channels conductivity, or a direct influence of *Reverbα* on L-type calcium channels to support day-time depolarisation of SCN clock cells (Jackson, 2004; Belle *et al*., 2009; Schmutz *et al*., 2014; Allen *et al*., 2017). Therefore, the delayed LGN neuronal activity phase relative to the *Per1* expression acrophase suggests potential diversity in the intracellular machinery connecting the TTFL to the membrane dynamics, or different ionic mechanisms coupling molecular and electrical clock.

Circadian oscillators may be divided into subclasses, according to the degree of autonomy in their rhythmic properties. The rhythm of a ‘slave oscillator’ is just a fingerprint of a rhythmic input from a stronger clock which damps and disappears in isolated conditions. This stands in contrast to ‘semi-autonomous’ oscillators which possess intrinsic timekeeping properties, however without the phasic input, its single cell rhythms desynchronise and the whole-field oscillation amplitude dampens quickly. Finally, the fully ‘autonomous’ clock persists robust single cell oscillations that are synchronised due to its internal connectivity (Begemann *et al*., 2020). In our study, the rhythmic clock gene expression of the DLG seen *in vivo* diminished in culture. Additionally, previous studies reported day-to-night change in the DLG neuronal activity *in vivo* (Brown *et al*., 2011), which are not present in our long-term *ex vivo* MEA recordings. Thus, it is tempting to classify the DLG as a ‘slave oscillator’ which requires an input from another distant brain clock to sustain its rhythmic properties. In contrast, both the IGL and VLG may be categorised as ‘semi-autonomous’ or ‘autonomous’ oscillators. Their rhythmic clock gene expression seen *in vivo* persists up to a week in culture and these molecular oscillations are also depicted in their well-organised circadian pattern of neuronal activity.

Owing to a strong input from the retinal ganglion cells, clock cells in the SCN respond to changes in ambient light. These acute electrical responses trigger molecular rhythms to shift; light appearing at the beginning of the dark phase delay behavioural activity rhythms, while the exposure to light at the end of activity phase is potent in phase advancing (Challet *et al*., 2003; Fuller *et al*., 2008; Takahashi, 2017). Here, we used optogenetic manipulation of retinal terminals in the LGN at projected ZT12 to recreate a phase delaying stimulus in the isolated slice condition without an input from the central clock. Neuronal activity recorded from control slices (i.e. these obtained from rats that were injected with the virus, but were not optogenetically stimulated during the recording) peaked earlier, compared to slices harvested from naïve rats. We speculate that the robust expression of *Chronos* opsin in the retina may have changed its phototransduction mechanisms or extended photoexcitability to the retinal ganglion cells beyond the melanopsin-expressing subset that are not intrinsically photosensitive. However, optogenetic stimulation of *Chronos*-expressing slices at the beginning of the projected night was potent to selectively delay activity peak (by ∼3 - 4.5 h) of the two compartments of VLG, without entraining the IGL. As the VLGm is generally not retinorecipient, or very sparsely innervated by the retina compared to the VLGl (Harrington, 1997; Beier *et al*., 2020), the phase shift in the VLGm is most likely to be transmitted via the VLGl. This is not surprising, as these two parts of VLG were demonstrated to stay in a strong reciprocal connection (Harrington, 1997; Govindaiah & Cox, 2009; Monavarfeshani *et al*., 2017). Intriguingly, the IGL phase was not affected by the optogenetic activation of retinal input. This suggests different non-photic Zeitgebers to possibly entrain its timekeeping, with the food intake or arousal being the most likely (Fernandez *et al*., 2020).

The IGL was classically theorised to transmit non-photic information to the SCN, via its NPY-synthesising neurons (Moore & Card, 1994; Harrington, 1997; Morin & Allen, 2006). These cells are not directly retinorecipient, but rather they gather light information from a second, directly retinorecipient subpopulation of IGL neurons which synthesise enkephalin (Thankachan & Rusak, 2005; Juhl *et al*., 2007; Blasiak & Lewandowski, 2013). Both of these IGL subpopulations are γ-aminobutyric acid (GABA)-ergic, with both NPY and enkephalin exerting inhibitory actions upon the IGL cells (Palus *et al*., 2017). Interestingly, our RNAscope protocol demonstrated that clock gene expression in the LGN does not co-localise with the mRNA of the enkephalin precursor (*Penk*). Thus, these directly retinorecipient IGL/VLG cells are not the likely candidate for circadian timekeeping neurons. In our MEA recordings we found more light-insensitive cells to change their firing rates in a circadian fashion, compared to these excited by optogenetic flashes. Hence, we form a hypothesis that light-sensitive IGL/VLG neurons are excited during the light phase *in vivo* and in turn inhibit clock cells, whose intrinsic neuronal activity rises during the night. This model differs from the light entrainment of the SCN, where clock cells are directly retinorecipient and thus more active during the day (Belle & Diekman, 2018; Hastings *et al*., 2019).

In summary, our study reveals previously unknown circadian timekeeping properties of the thalamic retinorecipient centre, the LGN. We propose the DLG to act as a slave oscillator, exhibiting rhythmic properties only when connected to its afferents *in vivo*. More importantly, we characterise novel intrinsic extra-SCN oscillators in the IGL and VLG, which organise their circadian physiology by means of autonomous rhythmic clock gene expression and circadian variability in their firing rate. We speculate that the higher night-time activity of the retinorecipient centres may help nocturnal animals to process limited ambient light information during their behaviourally active phase. Additionally, we find that the phase of the VLG neuronal activity is entrained by the optogenetic activation of retinal afferents to the LGN, thus we classify it as a light-entrainable oscillator – a feature shared with the SCN clock. This study provides a first description of local clock mechanisms at work in the LGN, and sets new directions for targeted physiological and pharmacological investigations of circadian biology of the visual system.

## Acknowledgements

We would like to thank the Department of Physiology and Toxicology of Reproduction for the access to their laboratory equipment. Authors would also like to thank Patrycjusz Nowik for the excellent animal care.

## Notes

**Funding**: This work was financially supported by a project ‘Sonatina 2’ 2018/28/C/NZ4/00099 given to LC and ‘Preludium 14’ 2017/27/N/NZ4/00785 to KP from the Polish National Science Centre. KP was additionally supported by ‘Etiuda 8’ doctoral scholarship 2020/36/T/NZ4/00341. JM was supported by the Taiwan Ministry of Science and Technology (109-2320-B-038-020, 109-2314-B-038- 071, 109-2314-B-038-106-MY3, 108-2321-B-006-023-MY2, 107-2410-H-038-004-MY2), the Higher Education Sprout Project by the Taiwan Ministry of Education (DP2-109-21121-01-N-01, DP2-110- 21121-01-N-01), and Taipei Medical University (TMU107-AE1-B15, 107-3805-003-110, 107TMU-SHH-03).

**Conflict of interest**: The authors declare no competing financial interests.

### Competing Interest Statement

The authors have declared no competing interest.

